# Physiotherapist confidence level in mobilising stroke patients after decompressive hemicraniectomy: are helmets useful?

**DOI:** 10.1101/632604

**Authors:** Sanjay Budhdeo, Toby Meek, Theodore D Cosco, Sanchit Turaga, Aswin Chari, Nikhil Sharma

## Abstract

**Introduction:** Decompressive hemicraniectomy is a lifesaving measure in malignant middle cerebral artery infarction; however, this leaves patients with a skull defect. There is variability of helmet use in this patient group across Britain. We aimed to examine whether (1) specialist physiotherapist were more confident mobilising a patient with hemiparesis and skull defect than a non-specialist physiotherapist (2) non-specialist and specialist physiotherapists would be more comfortable mobilising this patient with a helmet as opposed to without a helmet.

**Methods:** We carried out a cross-sectional online survey of specialist physiotherapists and non-specialist physiotherapists in Britain. Recruitment was through mailing lists. Physiotherapists were asked to rank their confidence level on a 5-point Likert scale of mobilising an example patient with and without a helmet. They were also asked about the number of additional therapists needed to safely mobilise the patient.

**Findings:** 96 physiotherapists completed the survey; 44 were specialists and 52 were non-specialists. Specialist physiotherapists felt more comfortable mobilising patients (mean difference = 0.68, p < 0.001). Non-specialist physiotherapists felt significantly more comfortable mobilising patients with a helmet (mean difference = 0.96, p value < 0.001), as did specialist physiotherapists (mean difference = 0.68, p value < 0.001). There was no difference in confidence level arising from helmet use between the two groups (p = 0.72).

**Conclusions:** Use of helmets may allow specialist and non-specialist physiotherapists to feel more comfortable when mobilising stroke patients post-decompressive hemicraniectomy. Consideration should be made by hospitals and health systems for the provision of helmets this patient group, to maximise functional gains.

## Introduction

### Hemicraniectomy reduces mortality in stroke

Large space occupying (or malignant) middle cerebral artery (MCA) infarction or hemispheric infarction represents approximately 1-10% of all strokes and has a grave prognosis (1). Without treatment, up to three quarters of these patients will die due to brain herniation (1). Early decompressive hemicraniectomy has been shown to reduce mortality, with the number needed to treat (NNT) for survival being 2.4, albeit with substantial morbidity (2). The hemicraniectomy performed for malignant MCA infarction is large, leaving a clearly visible skull defect. It is unknown whether the presence of this skull defect may influence the delivery of rehabilitation in the months following stroke, prior to cranioplasty; importantly, this period after stroke and surgery is critical for rehabilitation. Given the benefits of this procedure to patients, understanding the potential barriers to effective rehabilitation is important to providing optimal patient care. (2) (3) (4) (5) (6).

### Gait problems in malignant MCA infarcts

Gait problems in MCA strokes arise from hemiparesis which typically results in severe restrictions of mobility, as demonstrated DESTINY, DECIMAL and HAMLET trials (7). These gait issues could plausibly contribute to a higher risk of head injuries in postsurgical phase for patients with decompressive hemicraniectomy, especially without appropriate physiotherapy and head protection.

### Direct complications of the skull defect remain unexamined

There is a paucity of evidence in the literature regarding adverse direct complications from the skull defect following decompressive hemicraniectomy (7). One case report describes a patient who died due to haematoma formation at the site of skull defect following a fall (8). Helmets are an option for cranial protection prior to cranioplasty. The authors’ experience is that there is a variation of practice of helmet use in decompressive hemicraniectomy in UK neurosciences centres. This was confirmed by an informal survey of 10 neurosciences centres, suggesting that 50% used helmets. Studies analysing the physics of blunt trauma impact using helmets have provided evidence of the potential protective effect of a helmet (9) (10). No trials have been conducted into the use of helmets in patients post-decompressive hemicraniectomy, and whether they might provide protection for users. Given the rarity of direct complications of skull deficit (8), carrying out such a trial may require large numbers.

### Physiotherapist attitudes to the skull defect may influence rehabilitation

In light of the paucity of evidence to assess utility of helmets and the difficulties obtaining such data, an alternative way of assessing the utility of helmets and justifying a future trial would be to look at beliefs regarding helmets of healthcare practitioners involved in rehabilitation, and assessing whether the use of helmets would change the extent of therapy that patients receive.

After decompressive hemicraniectomy, patients are managed with a multidisciplinary approach. The physiotherapists involved are typically defined by work in a specific practice setting such as in hospital (neurosurgical, neurological, stroke, or rehabilitation wards) or in the community. Initially patients receive physiotherapy on the ward immediately post-surgery however, they would continue to receive therapy on discharge in the community. Community based care has less access to specialists and therefore physiotherapists may feel less confident in dealing with large skull defects post hemicraniectomy due to injury potential. The World Federation of Neurorehabilitation (WFNR) and European Association for Neurorehabilitation (EANR) both recommend specialised education for immediate postoperative care on the ward and long-term neuro-rehabilitation in the community. (11)

In order to assess confidence level levels for physiotherapists, we produced an online survey, which we sent to physiotherapists throughout England. Here, we define confidence level as the extent to which physiotherapists feel they can safely mobilise patients in an inpatient environment, for the purpose of undertaking activities related to rehabilitation. We used two proxies for physiotherapist confidence level. The first was a five-point Likert scale that assessed physiotherapist levels of confidence level when mobilising an example patient. The second was an estimation of the number of additional members of the therapist team that the physiotherapist thought would be needed to mobilise the example patient. We asked physiotherapists to consider these scenarios with and without a helmet.

Our primary aim was to study the possible impact of helmets on rehabilitation after hemicraniectomy for malignant MCA infarct. Our secondary aim was to understand whether any impact applied to all physiotherapist groups, and whether non-specialist physiotherapists were more affected. We tested three hypotheses. First, we hypothesised that there would be a difference between confidence levels between specialist and non-specialist physiotherapists. This would be expressed by absolute differences in self-described confidence level and opinions of number of therapists required to mobilise the patient without a helmet. Second, we hypothesised that use of a helmet will increase the confidence level of physiotherapists in mobilising the patient. This would be expressed as differences in the change in self-described confidence level between the two conditions, and opinions of number of therapists required to mobilise the patient between the same patient wearing and helmet and not wearing a helmet. Finally, our third hypothesis was that non-specialist physiotherapists would be more likely to have increased levels of confidence level from helmet use than specialist neuroscience physiotherapists. Currently, practice in the UK does involve the use of helmets in this patient cohort. This study is important because it may provide evidence for the use of helmets in decompressive hemicraniectomy patients.

## Methods

### Ethical Approval

As per work employing similar methodologies (12) and in accordance with UCL Research Ethics Committee guidelines, this work is focused on service development and fulfils criteria for exemption.

### Study design

The study was a cross sectional survey of physiotherapists who were members of specialist societies in the UK e.g. ACPIN (Association of Chartered Physiotherapists Associated in Neurology) and Chartered society of Physiotherapy. Links were disseminated through mailing lists and participants chose to be part of the survey. The Chartered Society of Physiotherapists has 58,000 chartered physiotherapists, physiotherapy students and support workers.

### Data Collection

SurveyMonkey (https://www.surveymonkey.com) was used to build a survey for physiotherapists to collect information in order to test hypotheses. The survey consisted of an explanation of decompressive hemicraniectomy through description of the procedure, an axial CT imaging slice showing a patient pre- and post-decompressive hemicraniectomy and YouTube video demonstrating a three-dimensional view of the skull defect after decompressive hemicraniectomy (https://www.youtube.com/watch?v=DQPSfXxOYYo). An introductory descriptor was used to explain the context of the survey, the purpose of hemicraniectomies: this can be found in the appendix.

Survey participants were then shown a YouTube video of a patient with stroke who has a hemiparetic gait, walking with assistance (https://www.youtube.com/watch?v=ag5Qq46VOGU). The YouTube videos were used under the Creative Commons license. They were asked to make reference to this video when answering the questionnaire. The patient’s head was not viewed in the video, which aided anonymisation and meant that the video was not biased to the helmet or non-helmet condition. The owner of the patient video and CT scan were contacted and permission for use was granted.

Survey participants were asked to rate their confidence level mobilising the patient on a five-point scale with and without helmet. Survey participants were also asked how many additional therapists they would require to feel comfortable mobilising the patient with and without a helmet. In addition, information regarding years of experience and practice setting of survey participants was collected.

In order to recruit survey participants, the Association of Chartered Physiotherapists Interested in Neurology and Chartered Society of Physiotherapists were contacted. In addition, individual hospitals in the East of England and Greater London areas were contacted by email. They survey opened in January 2016 and results were collected in May 2016.

### Characteristics of Physiotherapists

In this study, specialist neurological physiotherapists were defined as those who currently work solely in a neurology, neurosurgery, stroke or neurorehabilitation hospital setting. Non-specialist physiotherapists may include physiotherapists with other specialties in teaching hospitals, physiotherapists with a more general case mix in district general hospitals, or physiotherapists with a general practice in the community or rehabilitation setting. Participants were also asked to declare their years of practice as falling within 0-2 years, 2-5 years, 5-10 years, or more than 10 years of practice.

For self-described confidence level in mobilizing the patient in the video contained within the survey, the Likert scale values were described as follows: very comfortable: 5, somewhat comfortable: 4, neither comfortable nor uncomfortable: 3, somewhat uncomfortable: 2, very uncomfortable: 1. For the number of additional therapists participants would require to feel comfortable mobilising the featured patient, survey options were: none, one, two, or at least three.

### Data Analysis

Data was analysed using Stata version 14 (Stata Corp., College Station, TX). The first hypothesis examined the differences in self-described confidence level and in the number of additional therapists required between specialist and non-specialist physiotherapists. A logistic regression was undertaken (p <0.05 was considered significant). The independent variable was physiotherapist professional status (neurological specialist or non-specialist) and the dependent variable was physiotherapist confidence level (measured using self-described confidence level and number of additional therapists needed for mobilisation). Years of experience of the physiotherapists were controlled for.

The second hypothesis examined whether use of a helmet would result in differences in confidence level for physiotherapists. Paired t tests examined for differences in the helmet versus no helmet condition. This analysis was carried out separately for specialist and non-specialist physiotherapists.

The third hypothesis was that non-specialist physiotherapists would be more likely to report changes in confidence level in mobilising the patient as a result of helmet use than specialist neuroscience physiotherapists. In order to test this hypothesis, the specialist and non-specialist physiotherapists were divided by whether they had an increased level of confidence level with a helmet (expressed by a difference between self-described confidence level, or by a difference in the number of additional therapists they felt were needed). This was the dependent variable in an ordered logistic regression model. The independent variable was whether the physiotherapist was a specialist neurological physiotherapist, or whether they were a non-specialist physiotherapist. The number of years of experience was included as a covariate.

## Results

Participant characteristics are described in Table 1. In order to be a physiotherapist in the UK, one must have a registration with the Health and Care Professions Council, for which a degree level physiotherapy qualification is required (usually 3 year undergraduate or a two year accelerated Masters). The proportions of the group with 0-2 years or at least 10 years of experience was similar. 27.3% of the specialist neurological physiotherapist group had 2-5 years of experience whereas 40.4% of the non-specialist group had 2-5 years of experience. 40.9% of the specialist neurological physiotherapist group had 5-10 years of practice, whereas 23.1% of the non-specialist group had 4-10 years of practice. Given the disparities in experience between the specialist and non-specialist groups, an experience variable has been included as a covariate in all the analyses undertaken.

**Table 1:**
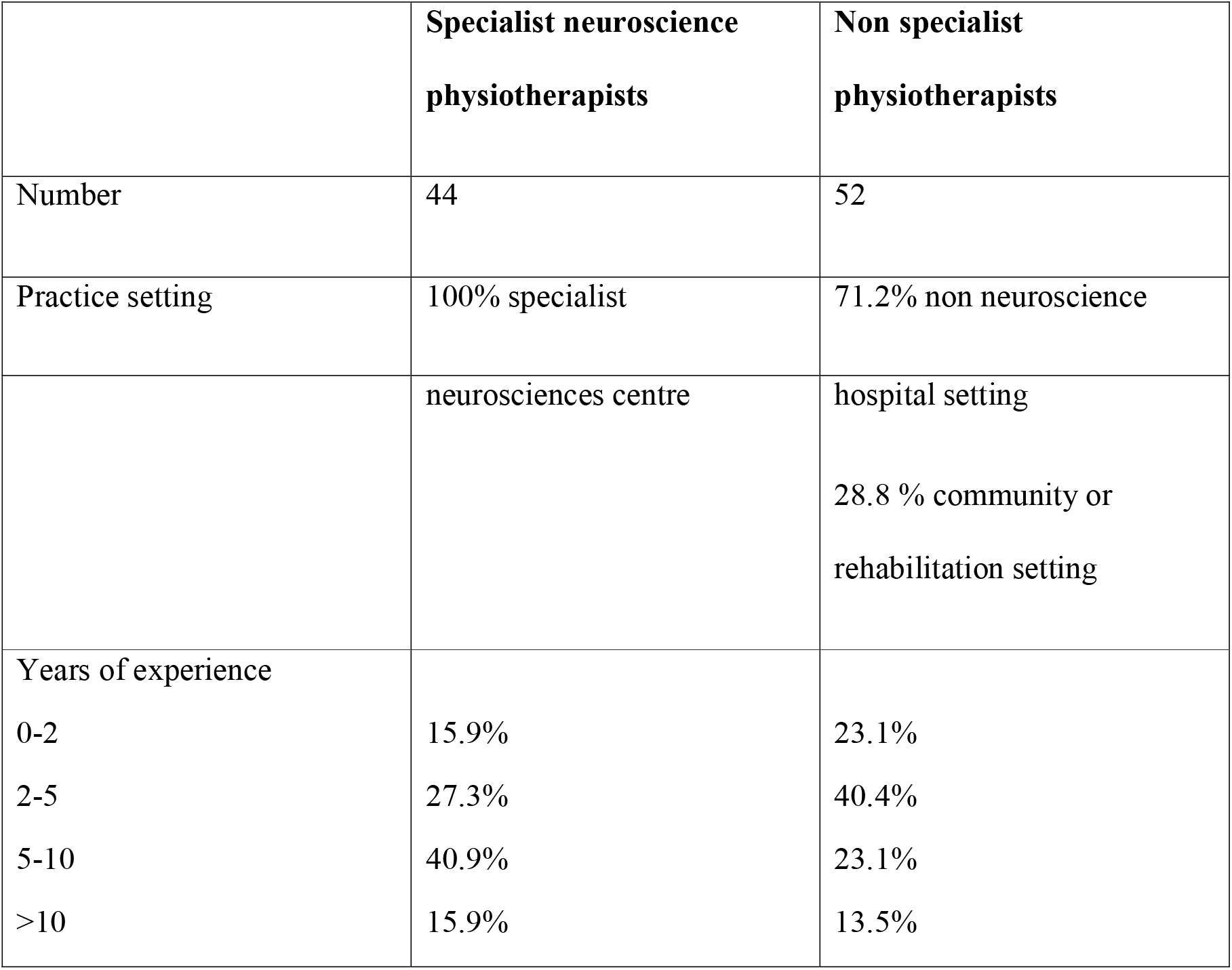
Participant characteristics of those who completed the survey.

|  | **Specialist neuroscience**<br>**physiotherapists** | **Non specialist**<br>**physiotherapists** |
| --- | --- | --- |
| Number | 44 | 52 |
| Practice setting | 100% specialist | 71.2% non neuroscience |
|  | neurosciences centre | hospital setting<br>28.8 % community or<br>rehabilitation setting |
| Years of experience |  |  |
| 0-2 | 15.9% | 23.1% |
| 2-5 | 27.3% | 40.4% |
| 5-10 | 40.9% | 23.1% |
| >10 | 15.9% | 13.5% |

When surveyed, specialist neurological physiotherapists report increased self-described confidence level mobilising stroke patients with decompressive hemicraniectomy than non-specialist physiotherapists in an experience-adjusted model (OR = 2.69, 95% CI 1.23-5.91 p value < 0.001). In contrast, there was no difference between the number of additional therapists that specialist neurological physiotherapists and non-specialist physiotherapists would prefer in an experience-adjusted model (OR = 0.70, 95% CI 0.31-1.58, p value = 0.15).

Non-specialist physiotherapists have increased self-described confidence level when mobilising stroke patients with decompressive hemicraniectomy if they are wearing helmets compared to no helmets (mean difference = 0.96, t value = 7.15, p value < 0.001). In addition, specialist neurological physiotherapists have increased self-described confidence level when mobilising stroke patients with decompressive hemicraniectomy if they are wearing helmets (mean difference = 0.68, t value = 3.51, p value < 0.001).

Non-specialist physiotherapists require fewer additional therapists when mobilising stroke patients with decompressive hemicraniectomy if they are wearing helmets (mean difference = −0.5, t value = 6.25, p value < 0.001). Specialist neurological physiotherapists require fewer additional therapists when mobilising stroke patients with decompressive hemicraniectomy if they are wearing helmets (mean difference = −0.41, t value = 5.45, p value < 0.001).

Examining the relationship between whether physiotherapist specialty and changing self-described confidence level depending on whether the patient was wearing a helmet, there was no significant association found (OR 0.86, 95% CI 0.38-1.97, p value = 0.72). Examining the relationship between whether physiotherapist specialty and changing the number of therapists required for assistance depending on whether the patient was wearing a helmet, there was no significant association found (OR 0.74, 95% CI 0.32-1.69, p value = 0.22). There was therefore no evidence that specialist neurological physiotherapists were less likely to exhibit a confidence for patients wearing a helmet than non-specialist physiotherapists.

## Discussion

Specialist physiotherapists were more comfortable mobilising stroke patients with decompressive hemicraniectomies; however, there was no evidence that they required a different number of additional therapists to aid with mobilisation. We also demonstrate that both specialist and nonspecialist physiotherapists would feel more comfortable and require fewer additional therapists to mobilise post-stroke patients with decompressive hemicraniectomy, were the patients to wear a helmet.

Our findings demonstrate that there is an association between increased physiotherapist confidence level mobilising patients and decompressive hemicraniectomy patients wearing helmets; however, there is no association between the additional the number of therapists required and wearing a helmet. This suggests that physiotherapist confidence level levels are intrinsic to patient state, rather than being associated with the amount of additional help available which is an important confounding factor. Relative staffing levels between hospitals cannot be implicated as a factor which might account for differences in therapy levels. Looking further at association between self-described confidence level and the helmet condition, the experience covariate is a significant confound. This suggests that more experienced physiotherapists feel more comfortable when working with this patient cohort, which would be expected given the complex nature of these patients, as regards impediments to mobility and safety.

In addition, we demonstrate that while specialist neurological physiotherapists are more comfortable mobilising stroke patients with decompressive hemicraniectomies, both specialist neurological physiotherapists and non-specialist physiotherapists feel more comfortable mobilising patients, were the patient to wear a helmet, providing a strong argument for future research into this area. This finding is interesting as it indicates that regardless of training and experience of this specialist area, physiotherapist change their attitudes to patients when they wear a helmet, and they regard the helmet as protective even if they are not given any evidence in support of this. Specialist physiotherapists appear to have a different relative threshold for mobilising patients, rather than different beliefs regarding suitability of mobilisation in this patient cohort. A further analysis (to explore whether there is a difference between specialty and non-speciality physiotherapists in how likely they were to change opinions on confidence level mobilising a patient between the helmet and no helmet condition) did not reveal a difference between the two groups. While we have made no judgements about the level of risk from mobilising stroke patients after decompressive hemicraniectomy, it is interesting to note that physiotherapist attitudes to whether helmets may be useful in mitigating risk of mobilisation do not differ with subject matter expertise

The helmet itself may present certain limitations in potential cost and aesthetic: helmets must be sufficiently light so as not to burden the patient but strong and stable enough to protect from head injury.

Helmets have been studied widely in many contexts where they have been shown to prevent head injury. In a study looking at cycle related injuries in those with helmets in 1040 patients, 114 of them wore helmets. Head injury was sustained by 4 people out of 114 (4%) as opposed to the higher proportion of 100 people out of 900 (11%). Moreover, odds ratios showed a protective factor of 3.25 (1.17 to 9.06, p=0.024) for wearing a helmet (21).

Helmets have been designed in the context of non-medical activities such as cycling but would likely need to be investigated and refined in the context of post-hemicraniectomy head injury. A study of 33 patients who had 14751 seizures in a one year period was conducted wherein they were provided with helmets. There were 59 injuries and helmets were only in use for 59% of accidents. In these situations, injuries continued to occur despite helmet use, particularly to the scalp and face (22). The study used ice hockey and hard foam (plastazote helmets), suggesting that a more refined approach specific to the nature of the potential injury is required. Indeed, many ice hockey helmets do not have facial protection and are often hard and heavy.

### Limitations

First, the survey was advertised in the UK, and respondents are from the UK. There is inter- and intra-country variability in the use of helmets after decompressive hemicraniectomy (13) (14) (15). Different countries may use helmets post hemi-craniectomy to different extents. Physiotherapists in different countries may have different attitudes towards the use of helmets. Second, the reach of the survey is unquantified; however, it is likely that only a small fraction of those who received the invitation to complete the survey responded to this request. There is a possibility of a biased sample due to this response rate. Third, this study is based on a video of one subject. The patient featured in the video used for this paper likely has a modified Rankin scale of 4, and this is typical for a patient who has had a malignant MCA infarct and decompressive hemicraniectomy (16). In any cohort of patients with decompressive hemicraniectomy following malignant MCA infarct, there will be variability of patient deficit in the immediate post-operative period and in the long term, so analysis of multiple patient videos representing differing levels of deficit would have improved the generalisability of this study.

Assessments using a Likert scale have limitations. An analysis of research into the Likert system showed that surveyed people tend to pick the central options more than extremes (i.e. very comfortable and very uncomfortable), termed the “anchor effect” (23). Furthermore, similarities between options such as “very comfortable” and “comfortable” may have had different meanings to different physiotherapists.

### Implications for practice

The estimated cost of stroke to the UK economy is £9 billion annually (17). Even small changes in functional ability can increase the independence of stroke patients (18) (19). If physiotherapists feel that patients are more able to partake in physiotherapy as a result of using a helmet, this may result in functional improvements in these patients. Reduced physiotherapy input because of safety concerns may unnecessarily limit treatment.

### Future directions of research

There has been insufficient study into the utility of helmets in decompressive hemicraniectomy patients. This is partly because adverse outcomes due to falls after hemicraniectomy are very rare. Some experts suggest use of a helmet in such circumstances (20), but this is not universal and practises vary between different centres and between different countries. Rather than seeking evidence for the efficacy of helmets in the setting of post-operative malignant middle cerebral artery patient, we adopt a novel approach. We examine whether there are benefits of helmets with regards to aiding physiotherapist mobilisation, rather than considering their intrinsic benefit.

Future work should address the lack of systematic study into the adverse consequences of mobilisation in decompressive hemicraniectomy patients. Even for large centres of excellence, there may not be sufficient cases for a case series. One option would be to set up a registry for post-hemicraniectomy complications related to mobilisation. Interventional studies could be carried out to assess whether helmets did result in improved functional outcomes for this patient cohort. An unblinded study could be straightforward to arrange, especially if randomisation occurred at the hospital level. Qualitative studies, such as semi-structured interviews, would be useful to explore the determinants of physiotherapist attitudes to mobilisation of post-stroke hemicraniectomy patients. Given the differences in helmet use internationally, comparison of physiotherapist responses across different countries may be particularly useful.

### Conclusion

Use of helmets increase physiotherapist confidence level immobilising stroke patients with decompressive hemicraniectomy. This is important because physiotherapy because the brain enters a heightened period of plasticity for a limited time post-stroke, and physiotherapy can be maximised during this period to improve patient outcomes.

## Authorship

SB planned the project. TM completed data collection. TDC completed data analysis. The first draft was written by SB. This was revised by TM, TDC, ST, AC and NS.

## Appendix 1 Descriptor used in the survey

“This survey is about patients with stroke who have had to undergo operations called decompressive hemicraniectomies. Large strokes can cause brain swelling. Brain swelling can cause death due to compression of the brainstem. Decompressive hemicraniectomy is a surgical technique used to relieve the increased pressure caused by the brain swelling and involves the removal of skull and an associated underlying layer of restrictive tissue covering the brain”

## Acknowledgements

None.

